# The anti-SARS-CoV-2 BNT162b2 vaccine suppresses mithramycin-induced erythroid differentiation and expression of embryo-fetal globin genes in human erythroleukemia K562 cells

**DOI:** 10.1101/2023.09.07.556634

**Authors:** Matteo Zurlo, Jessica Gasparello, Marco Verona, Chiara Papi, Lucia Carmela Cosenza, Alessia Finotti, Giovanni Marzaro, Roberto Gambari

## Abstract

The COVID-19 severe acute respiratory syndrome coronavirus 2 (SARS-CoV-2) the ongoing coronavirus disease 2019 (COVID-19) pandemic. The SARS-CoV-2 Spike protein (S-protein) plays an important role in the early phase of SARS-CoV2 infection through efficient interaction with ACE2. The S-protein is produced by RNA-based COVID-19 vaccines, and has been hypothesized to be responsible for damaging cells of several tissues and for some important side effects of RNA-based COVID-19 vaccines. The aim of this study was to verify the effect of the BNT162b2 vaccine on erythroid differentiation of the human K562 cell line, that has been in the past intensively studied as a model system mimicking some steps of erythropoiesis. We found that the BNT162b2 vaccine suppresses mithramycin-induced erythroid differentiation of K562 cells. Reverse-transcription-PCR and Western blotting assays demonstrated that suppression of erythroid differentiation was associated with sharp inhibition of the expression of α-globin and γ-globin mRNA accumulation. Inhibition of accumulation of ζ-globin and ε-globin mRNAs was also observed. In addition, we provide in silico studies suggesting a direct interaction between SARS-CoV-2 Spike protein and Hb Portland, that is the major hemoglobin produced by K562 cells. This study thus provides information suggesting the need of great attention on possible alteration of hematopoietic parameters following SARS-CoV-2 infection and/or COVID-19 vaccination.

## 1. Introduction

The coronavirus disease 2019 (COVID-19) pandemic has represented one of the major health problems since 2020 [1–4]. The fight against the COVID-19 severe acute respiratory syndrome coronavirus 2 (SARS-CoV-2) has been effective thanks to extensive vaccination campaigns that have been made possible by the approval of anti-SARS-CoV-2 vaccines by the Regulatory Agencies [4–10]. Starting from these approvals, COVID-19 vaccines have been extensively tested and distributed worldwide [4–6]. Thanks to the extensive use of COVID-19 vaccines and the improvement of the management of COVID-19 patients, the pandemic is at present under control, as stated by the recently issued WHO position (May 5 2023), concurring with the advice offered by the Report of the fifteenth meeting of the International Health Regulations (IHR) Emergency Committee. In this Report it was determined that that COVID-19 is now an established and ongoing health issue which no longer constitutes a public health emergency of international concern [11]. Accordingly, it has been suggested that it is time for further and more extensive evaluations of the short- and long-term effects of the COVID-19 vaccines on human tissue systems [12].

Concerning this very important issue, studies have hypothesized that the SARS-CoV-2 Spike protein (S-protein) is a major factor accounting for side effects of the COVID-19 mRNA vaccines [7–13], such as the BNT162b2 from Pfizer-BioNTech [14] and the mRNA-1273 from Moderna [15], as recently discussed by Trougakos and colleagues [16,17].

Relevant to this issue are reported evidences of circulating Spike Protein detected in post-COVID-19 mRNA vaccinated subjects [18–23], and several reports outlining that S-protein affects cellular metabolism and gene expression in a variety of tissue systems [12, 24–29], including the hematopoietic system [28,29]. Accordingly, the effects of SARS-CoV-2 infection and/or vaccination on the hematopoietic system should be carefully considered [28–30].

The objective of this study was to determine whether an an-ti-SARS-CoV-2 RNA vaccine (the BNT162b2) on the well-known K562 cell line [31], that has been in the past intensively studied as a model system mimicking some steps of erythropoiesis [32–34]. K562 cells express baseline levels of globin genes, the most important being those coding the embryo-fetal α-, ζ-, ε- and γ-globins. HbPortland ζ_2_γ_2_ is the predominant hemoglobin produced by K562 cells. Hb Gower 1 (ζ_2_ε_2_) and fetal hemoglobin (HbF, α_2_γ_2_) are also produced. In addition, the expression of globin genes and the progression through the erythroid differentiation pathway can be induced by a variety of HbF inducers [35–37]. Accordingly, K562 cells have been used not only as a model system to study the regulation of the expression of embryo-fetal globin genes [38], but also as a model system for the screening of inducers of HbF, of potential interest in the therapy of β-thalassemia and sickle-cell disease (SCD) [37]. In fact, it is well established that HbF production is beneficial for both β-thalassemia and SCD [39].

Another important point, relevant in respect to the focus of our study, is that K562 cells treated with anti-SARS-CoV-2 Comirnaty (BNT162b2) and Spikevax (mRNA-1273) COVID-19 vaccines, produce and release high levels of the SARS-CoV-2 S-protein, encoded by the mRNAs delivered by the vaccine formulations [40].

In this study, the effects of the Pfizer-BioNTech BNT162b2 vaccine were analyzed on erythroid differentiation and expression of globin genes in K562 cells cultured in the absence or in the presence of the HbF inducer mithramycin (MTH) [41,42]. Accumulation of globin mRNA was studied by RT-qPCR and globin and hemoglobin production by Western blotting and by benzidine staining of the treated K562 cells [41]. Mithramycin was selected for most of the experiments here reported, considering the fact that is one of the most potent erythroid inducers of K562 cells [41,42].

## 2. Results

### 2.1. Effect of the COVID-19 BNT162b2 vaccine on proliferation of K562 cells

Figure 1A shows that treatment of K562 cells with increasing concentrations of the BNT162b2 vaccine causes a dose-dependent inhibition of K562 cell growth. The data presented in Figure 1 indicate that 0.5 μg/mL of COVID-19 BNT162b2 vaccine was sufficient to cause inhibition of cell growth of treated K562 cells. This was found highly reproducible and the maximum effect being obtained with 2 μg/mL BNT162b2 concentration (p < 0.01). As expected, the intracellular content of SARS-CoV-2 Spike protein mRNA increases depending on the employed concentrations (Figure 1B). A significant increase (p < 0.01) was observed when 1 μg/mL vaccine was used.

**Figure 1.**
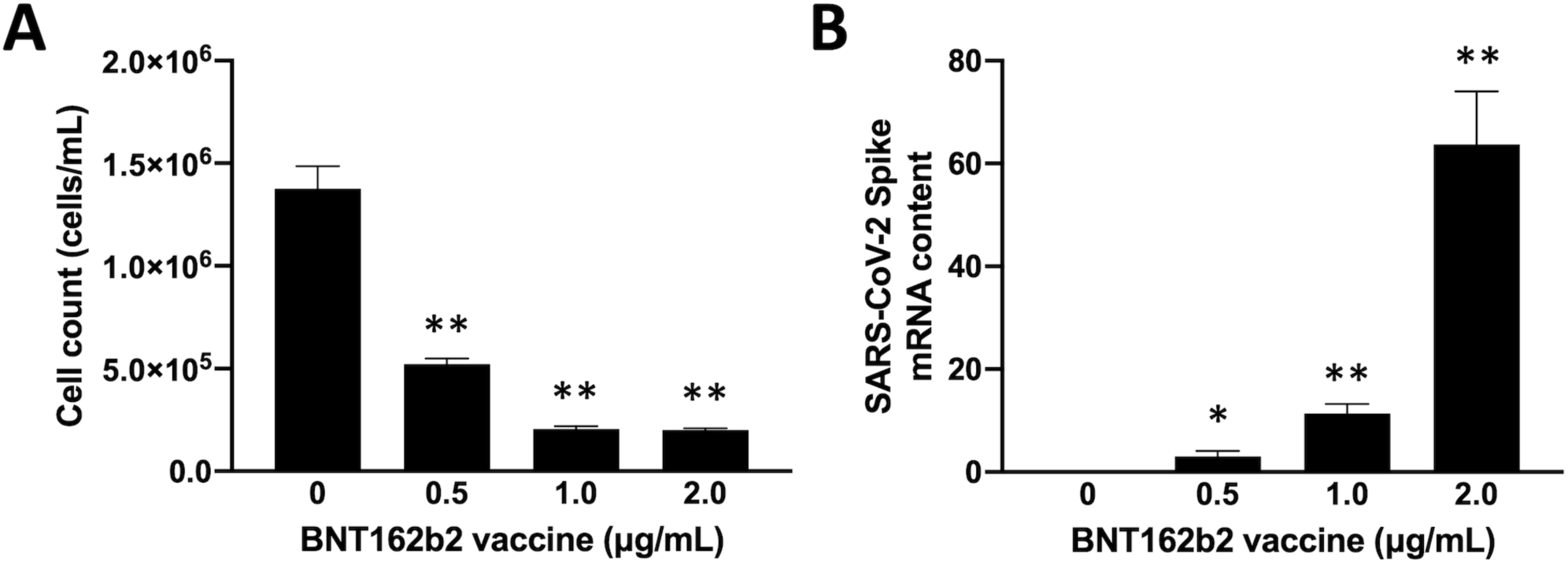
Inhibitory effects of COVID-19 BNT162b2 vaccine on cell proliferation of K562 cells. A. Cells were treated with the indicated concentrations of BNT162b2 vaccine and cell number/mL determined after 6 days of cell growth. B. Content of SARS-CoV-2 Spike mRNA sequences in BNT162b2-treated cells. Results are presented as mean ± S.E.M; statistical differences between groups were compared using ANOVA. (*): p < 0.05 (significant); (**): p < 0.01 (highly significant).

As expected from previously published observations [40], and in agreement with Figure 1A, production of S-protein was detectable when Western blotting was performed using cellular lysates from K562 cells treated with the BNT162b2 vaccine (Supplementary Figure S1). As expected from the notion that in many cellular systems the S-protein induces the expression of pro-inflammatory genes through up-regulation of NF-kB [26,43,44], increase of expression of NF-kB was found in K562 cells treated with the BNT-162b2 vaccine Supplementary Figure S2).

### 2.2. Inhibitory effect of the BNT162b2 vaccine on constitutive expression of globin genes in treated K562 cells

Figure 2 shows that treatment of K562 cells with the BNT162b2 vaccine causes a dose-dependent inhibition of the intracellular content of ζ-globin, α-globin, ε-globin and γ-globin mRNAs. The expression of β-globin gene was not assessed, as these gene is not expressed by K562 cells, that are on the contrary committed to high expression of embryo-fetal globin genes as reported in several studies [36,37]. When cells are exposed to the BNT162b2 vaccine, full inhibition of expression of α-globin (Figure 2A), γ-globin (Figure 2B), ε-globin (Figure 2C) and ζ-globin (Figure 2D) genes was obtained and correlates with a sharp enhancement of Spike mRNA content (Figure 1B). The response to the low BNT162b2 concentration (0.5 μg/mL) is to some extent surprising, since indicates and apparent increase of globin gene expression. This should be further studied and might be due to the non-RNA constituent(s) of the BNT162b2 vaccine.

**Figure 2.**
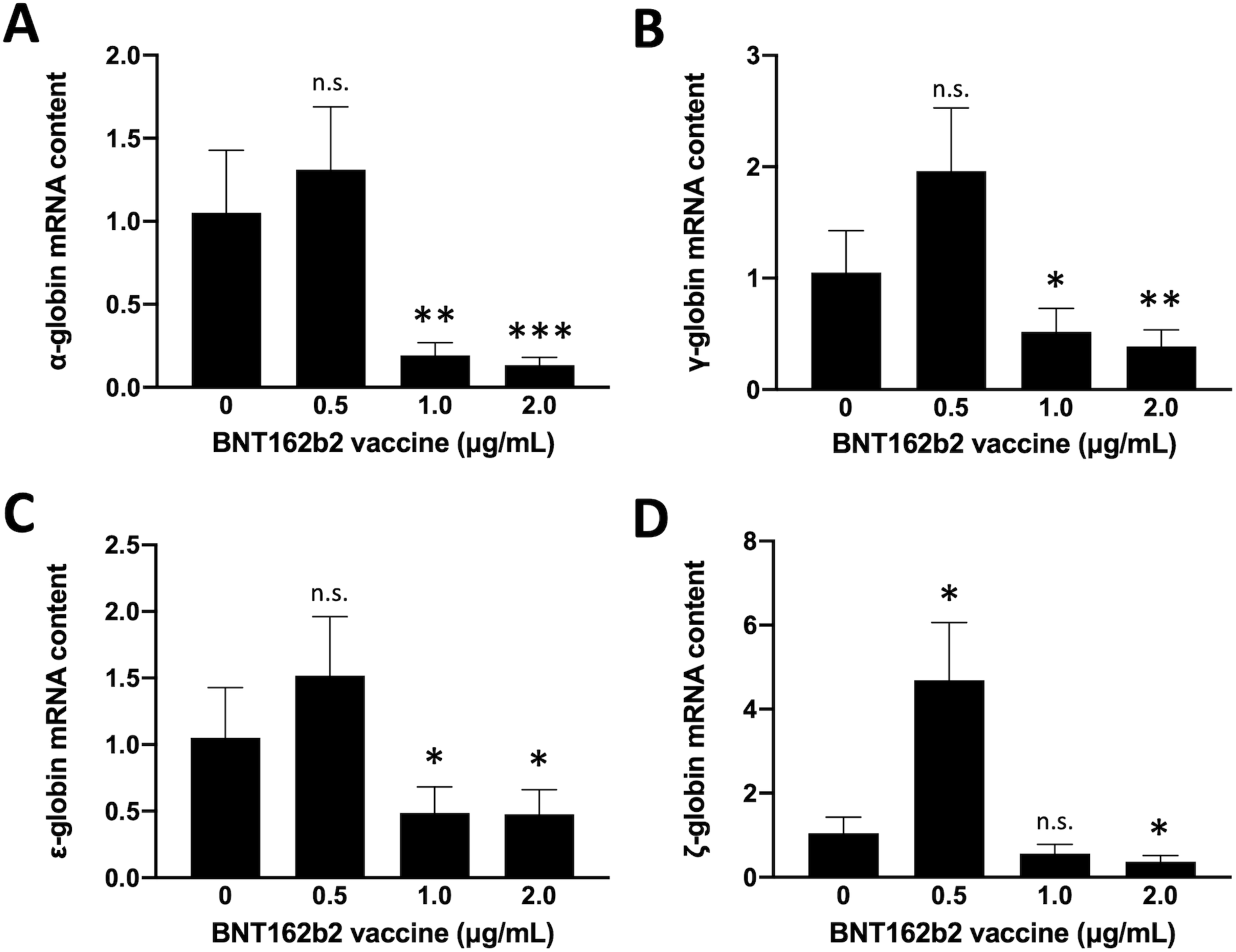
Effects of BNT162b2 vaccine on constitutive expression of embryo-fetal globin genes in K562 cells. Cells were treated in the absence or in the presence of the indicated amounts of BNT162b2. After 6 days RNA was isolated and RT-qPCR performed to quantify α-globin (A), γ-globin (B), ε-globin (C) and ζ-globin (D) mRNAs. Results are presented as mean ± S.E.M; statistical differences between groups were compared using ANOVA. (n.s.): not significant; (*): p < 0.05 (significant); (**): p < 0.01 (highly significant); (***): p < 0.001 (highly significant).

### 2.3. Treatment of K562 cells with COVID-19 BNT162b2 vaccine suppresses mithramycin induced erythroid differentiation

One of the most studied biological properties of the K562 cell line is that it can undergo erythroid differentiation upon exposure to a large variety of chemical inducers such as hemin, hydroxyurea, mithramycin, butyric acids and analogues, rapamycin, resveratrol and many others [35–37]. One of the most powerful compounds is the DNA-binding drug mithramycin (MTH). Erythroid differentiation can be assayed by the simple benzidine test, that is able to mark hemoglobin production. While uninduced K562 display a very low proportion of benzidine (hemoglobin-containing) cells (usually not exceeding 5%), when they are cultured with mithramycin the proportion of benzidine-positive cells sharply increases to 60-70% after 4-5 days of cell culture, as first reported by Bianchi et al. [41]. Figure 3 shows that treatment of K562 cells with the BNT162b2 vaccine causes a dose-dependent inhibition of erythroid differentiation. In panels A-D of Figure 3, benzidine staining of MTH treated K562 cells (Figure 3A) is compared to that of K562 cells treated with 0.5, 1 and 2 μg/mL of the BNT162b2 vaccine (Figure 3B, C and D). In Figure 3H the kinetic of erythroid differentiation is shown, demonstrating the inhibitory effects of BNT162b2. This inhibitory effect is highly reproducible, as indicated by Figure 3I.

**Figure 3.**
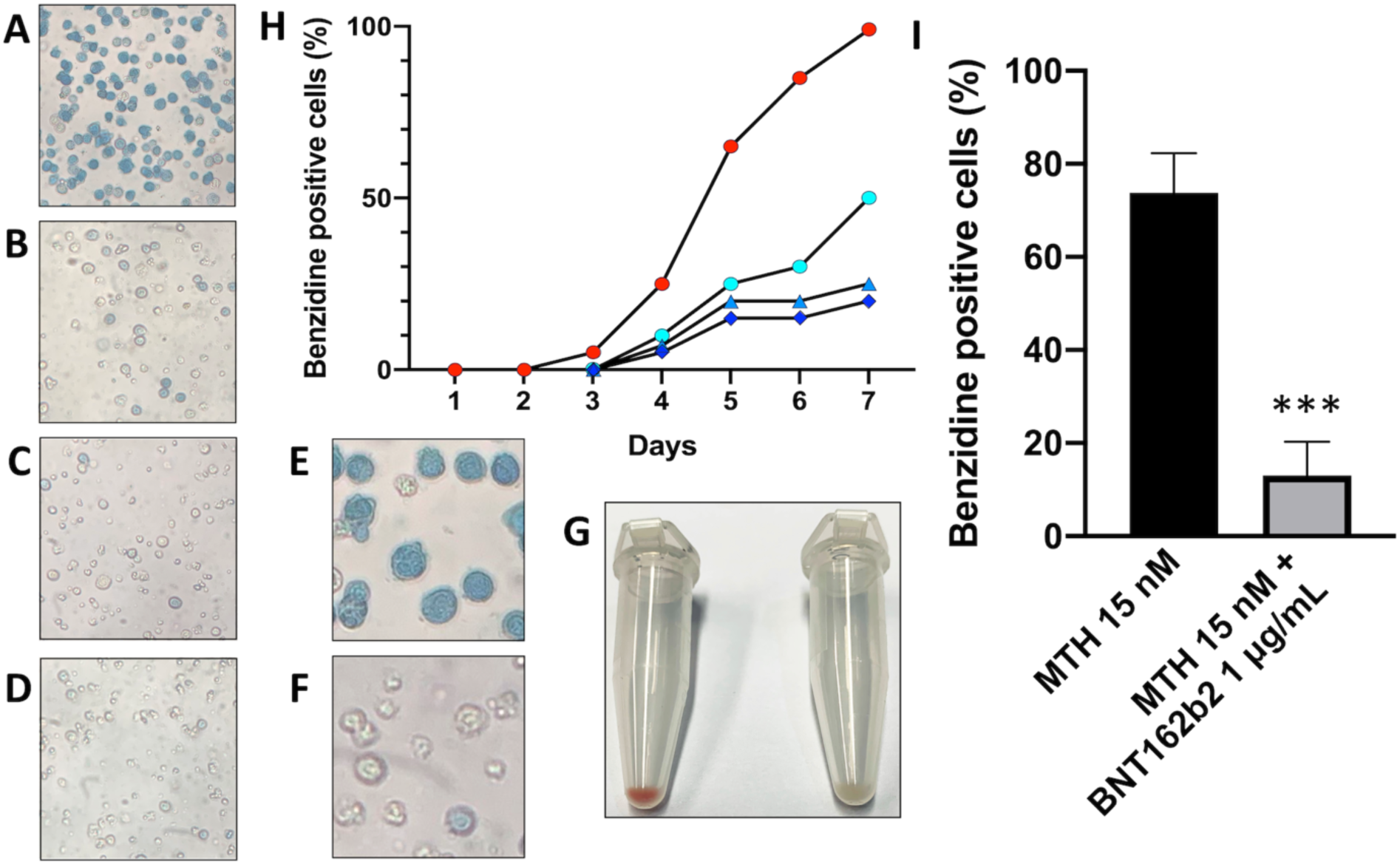
Effects of BNT162b2 vaccine on erythroid differentiation of K562 cells evaluated by the benzidine staining. A-D. K562 cells cultured for 5 days in the presence of 15 nM MTH (A) or MTH and 0.5, 1 and 2 μg/mL of BNT162b2 (B, C, D), magnitude 20x. E-F. Particular of image A and D respectively, showing the difference in hemoglobin production by benzidine staining at magnitude 40x. G. Comparison of cellular pellet obtained by centrifugation of K562 cells treated with 15 nM MTH (on the left) and MTH in the presence of 1 μg/mL of BNT162b2 vaccine (on the right). H. Kinetic of the increase of the % of benzidine-positive cells in K562 cells treated with 15 nM MTH (red circles), with 15 nM MTH and 0.5 μg/mL of BNT162b2 (azure open circles), with 15 nM MTH and 1 μg/mL of BNT162b2 (blue triangles), or with 15 nM MTH and 2 μg/mL of BNT162b2 (blue squares). I. Summary of 4 independent experiments comparing K562 cells induced with 15 nM MTH (black histogram) to cells induced with MTH in the presence of 1 μg/mL BNT162b2 (grey histogram) after 5 days of treatment. Results in panel I are presented as mean ± S.E.M; statistical differences between groups were compared using paired t-test. (***): p < 0.001 (highly significant).

In conclusion, these set of data demonstrate that the BNT162b2 vaccine suppresses mithramycin-induced erythroid differentiation of K562 cells.

### 2.4. The COVID-19 BNT162b2 suppresses erythroid differentiation in K562 cells treated with different inducers

As already pointed out, one of the most studied biological properties of the K562 cell line is that it undergoes erythroid differentiation upon exposure to a large variety of chemical inducers [35–37]. In order to determine whether the inhibitory effect of BNT162b2 is reproducible using other inducers of K562 erythroid differentiation, we employed the following inducers: rapamycin [45], hydroxyurea [46], resveratrol [47] and the isoxazole analogue c4 [48]. K562 cells were induced with 200 nM rapamycin, 200 mM hydroxyurea, 30 μM resveratrol, 150 nM c4 and 15 nM mithramycin (used as reference positive control) in the absence or in the presence of 1 μg/mL of BNT162b2. After 4 and 5 days, cells were harvested and analyzed with the benzidine assay, to detect hemoglobin producing cells.

The results obtained are reported in Figure 4 and clearly indicate that treatment of K562 with the BNT162b2 vaccine suppresses hemoglobin accumulation irrespectively of the employed inducers. More information on this experiment are shown in Supplementary Figure S3.

**Figure 4.**
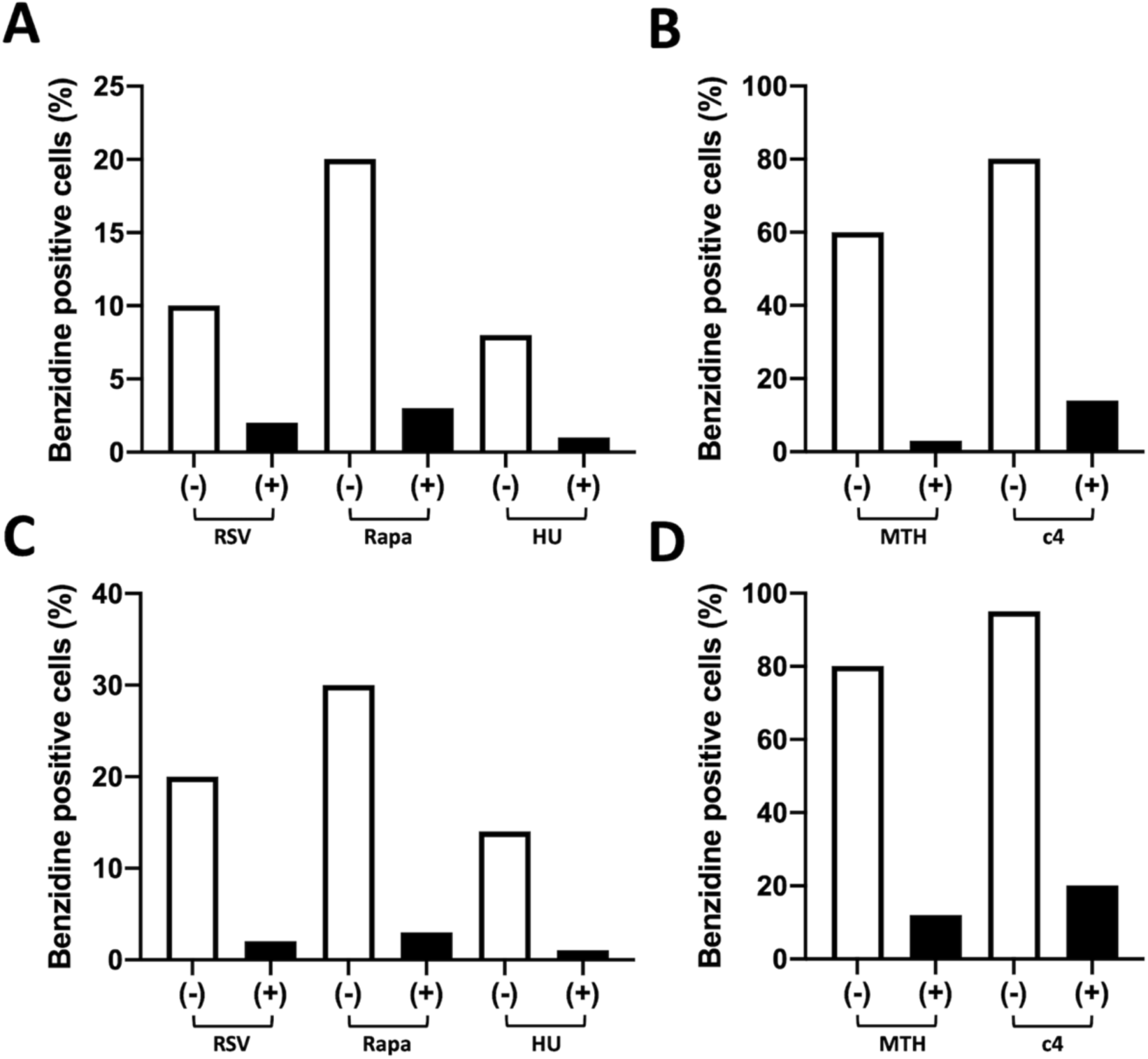
Effects of BNT162b2 vaccine on erythroid differentiation induced in K562 cells by different inducers. K562 cells were induced with 30 μM resveratrol (RSV) (A,C), 200 nM rapamycin (Rapa) (A,C), 200 μM hydroxyurea (HU) (A,C), 150 nM isoxazole c4 (B,D) in the absence (-) or in the presence (+) of 1 μg/mL BNR162b2, as indicated. After 4 (A,B) and 5 (C, D) days benzidine assay was performed. 15 nM Mithramycin (MTH) (B,D) was used as a positive control.

### 2.5. Inhibitory effect of the BNT162b2 vaccine on MTH-induced expression of globin genes in treated K562 cells

Figure 5 shows that treatment of K562 cells with the BNT162b2 vaccine causes a dose-dependent inhibition of the intracellular content of MTH-induced ζ-globin, α-globin, ε-globin and γ-globin mRNAs. First, the uptake of SARS-CoV-2 S protein mRNA was very efficient in MTH-induced, BNT162b2 treated K562 cells. This is depicted in Figure 5A, that clearly shows that, as expected, high levels of S-protein mRNA Sequence are detectable in cells treated with the BNT162b2 vaccine. The increase of the content of S-protein mRNA becomes highly significant when cells are treated with 1 and 2 μg/mL BNT162b2.

**Figure 5.**
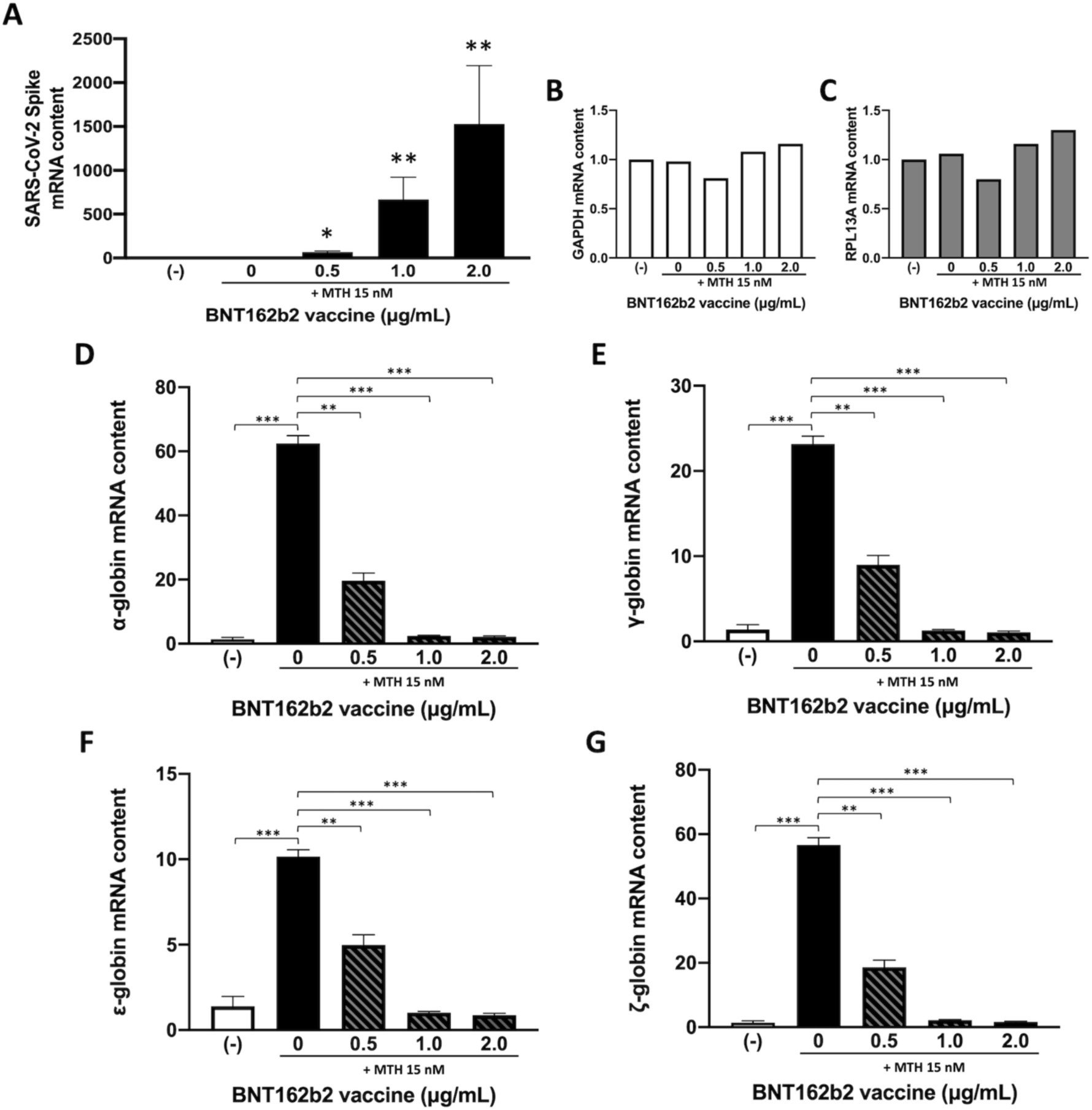
Effects of BNT162b2 vaccine on MTH-induced expression of embryo-fetal globin genes. After 5 days of K562 cell culturing as indicated, RNA was isolated and RT-qPCR performed to quantify the mRNA coding for: SARS-CoV-2 S protein (A), GAPDH (B), RPL13A (C), α-globin (D), γ-globin (E), ε-globin (F) and ζ-globin (G). Results are presented as mean ± S.E.M; statistical differences between groups were compared using ANOVA. (*): p < 0.05 (significant); (**): p < 0.01 (highly significant); (***): p < 0.001 (highly significant).

In this set of experiments, MTH induction was very effective. In fact, the fold increase of globin mRNAs ranged from 10.15±0.4 for ε-globin mRNA (Figure 5F) to 62.42±2.44 for α-globin mRNA (Figure 5D). In the presence of the BNT162b2 vaccine the content of all the studied mRNAs was found dramatically lower, indicating a suppression of the MTH-induction of globin gene expression in BNT162b2-treated cells. All these analyses were performed using β-actin as endogenous control housekeeping gene sequence. The % BNT162b2-mediated inhibition of the expression of ζ-globin, α-globin, ε-globin and γ-globin genes, was 67.2%, 68.4%, 50.1% and 60.7%, respectively (considering 0.5 μg/mL BNT162b2 concentration). When the 1 μg/mL BNT162b2 concentration was employed, the % inhibition values obtained were 96.1% (ζ-globin mRNA), 96.2% (α-globin mRNA), 90.1% (ε-globin mRNA) and 94.5% (γ-globin mRNA) (Figure 5). The effect of BNT162b2 is detectable even when the lowest BNT162b2 concentration was considered (0.5 μg/mL). On the contrary, the expression of two housekeeping genes (GAPDH and RPL13A) was unaffected (Figure 5, panels B and C).

### 2.6. The BNT162b2 vaccine inhibit the endogenous and MTH-induced accumulation of embryo-fetal globins

In consideration of the importance of γ-globin production for the severity of hematopoietic diseases (such as β-thalassemia and Sickle-cell disease), we determined whether treatment of K562 cells with the BNT162b2 vaccine is associated with inhibition of globin chains production at protein level. To this aim untreated and MTH-treated K562 cells were cultured in the presence of 0.5, 1 and 2 μg/mL BNT162b2 for 3 days, proteins were isolated and Western blotting performed to detect the production of the different embryo-fetal hemoglobin chains. Figure 6 shows that treatment of K562 cells with the BNT162b2 vaccine causes a dose-dependent inhibition of the intracellular content of all globin chains in both untreated and MTH-treated K562 cells. These data are fully in agreement with those presented in Figure 2 and Figure 5 and demonstrate that the BNT162b2 vaccine strongly inhibits accumulation of embryo-fetal globins (Figure 6).

**Figure 6.**
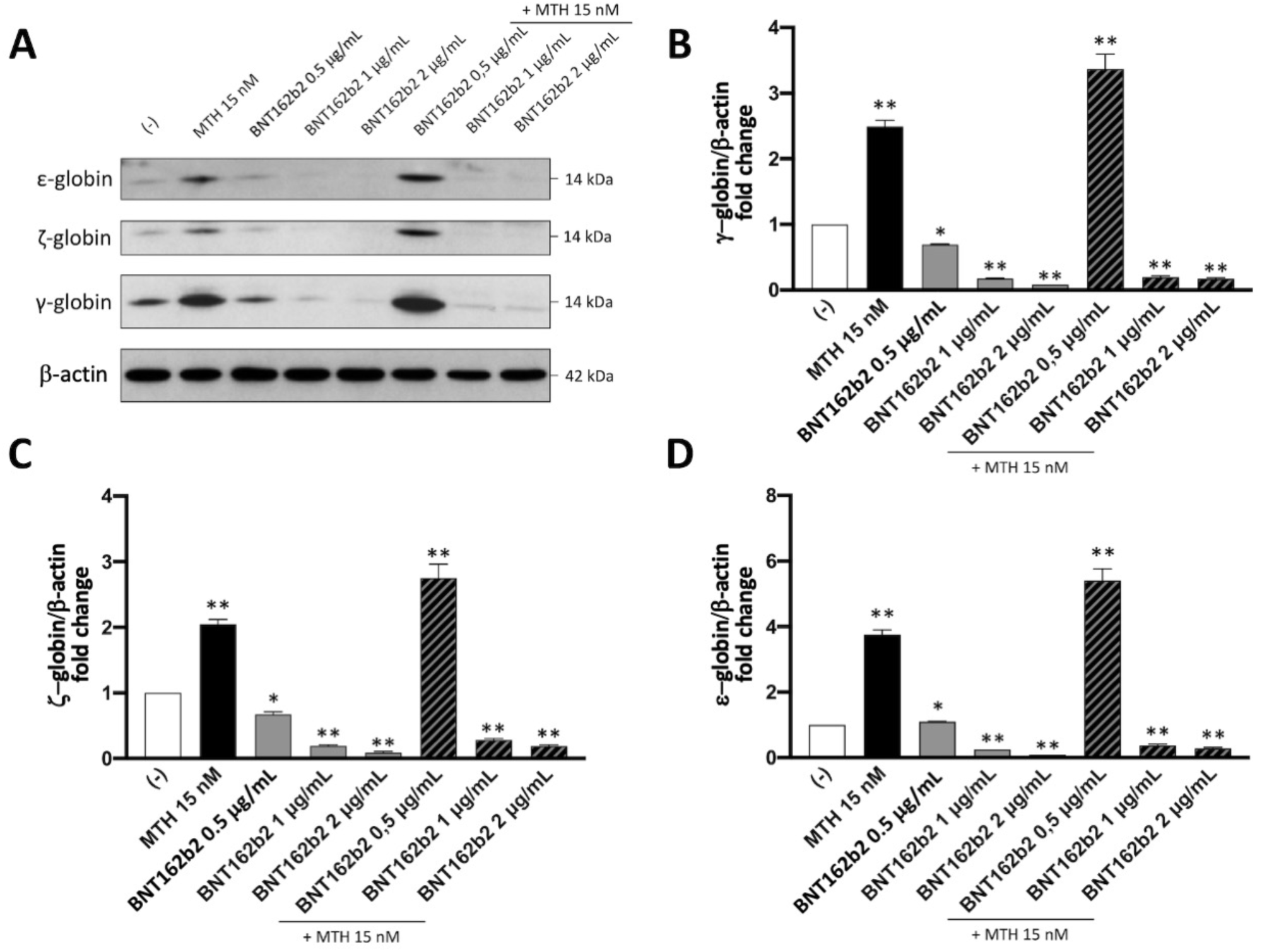
Effects of BNT162b2 on accumulation of γ, ζ, and ε-globin chains. K562 cells were treated as indicated for 5 days, then isolated proteins were analyzed by western blotting using rabbit anti-γ-globin (PA5-29006, Thermo Fisher Scientific Inc., Waltham, MA, USA), rabbit anti-ζ (Α6920, ABclonal, Woburn, MA, USA) and anti-ε globin (Α3909, ABclonal, Woburn, MA, USA) as primary antibodies, specific for the human γ, ζ, and ε-globin chains A. Representative image of obtained Western blot; B. densitometry analysis. Results are presented as mean ± S.E.M; statistical differences between groups were compared using ANOVA. (*): p < 0.05 (significant); (**): p < 0.01 (highly significant).

Interestingly, the expression of other genes involved in erythroid differentiation, such as the transferrin receptor gene was not affected by BNT162b2 treatment (Figure 7), suggesting that the inhibition of the expression of globin genes in K562 cells treated with BNT162b2 is specific.

**Figure 7.**
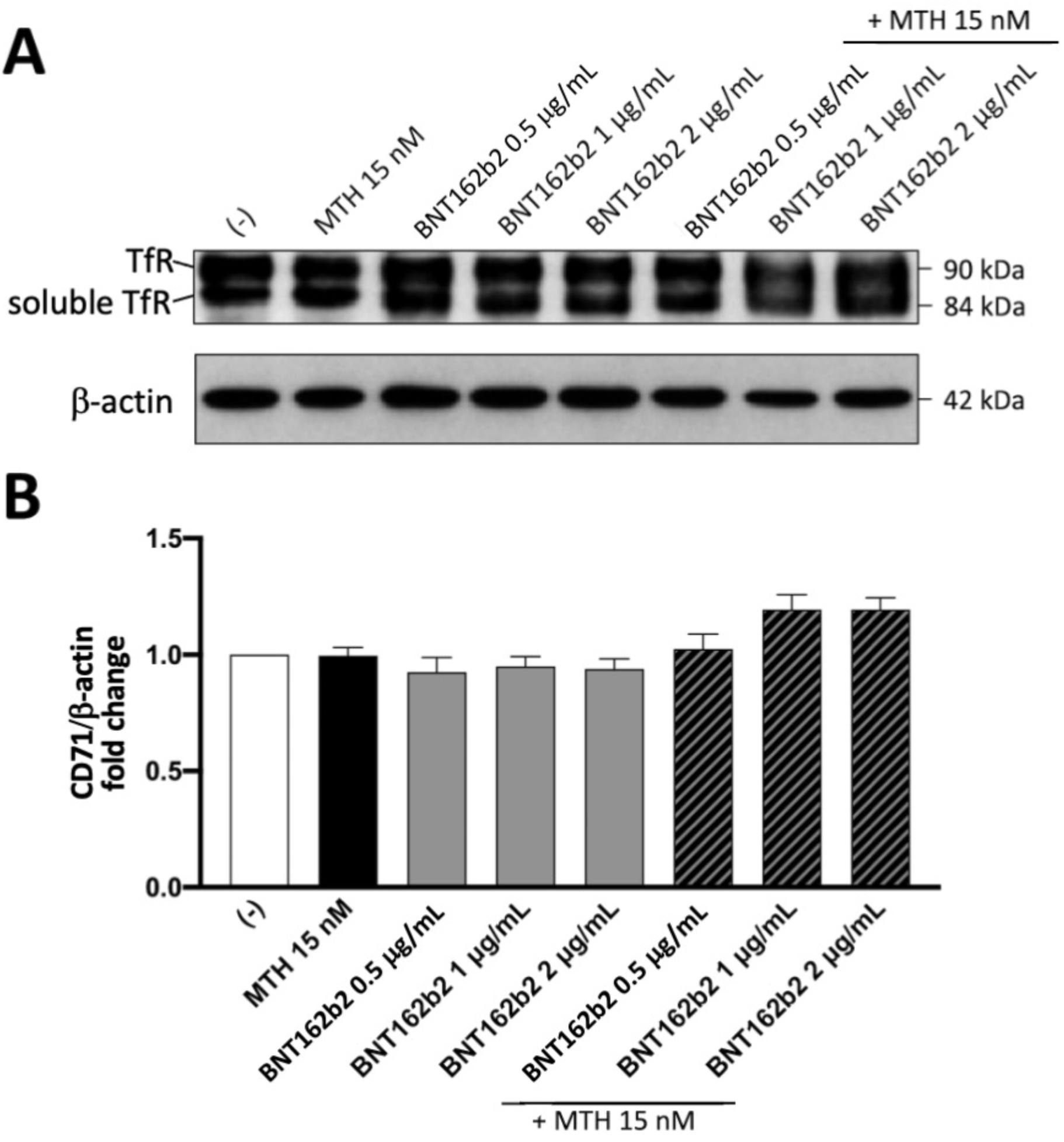
Effects of BNT162b2 on accumulation of transferrin receptor. K562 cells were treated as indicated for 6 days, then isolated proteins were analyzed by western blotting using anti-CD71 rabbit primary antibody (A22161, Abclonal, Woburn, MA, USA) specific for the human transferrin receptor. A. Representative image of the Western blotting obtained; B. densitometry analysis. Results are presented as mean ± S.E.M; statistical differences between groups were compared using ANOVA.

**Figure 8.**
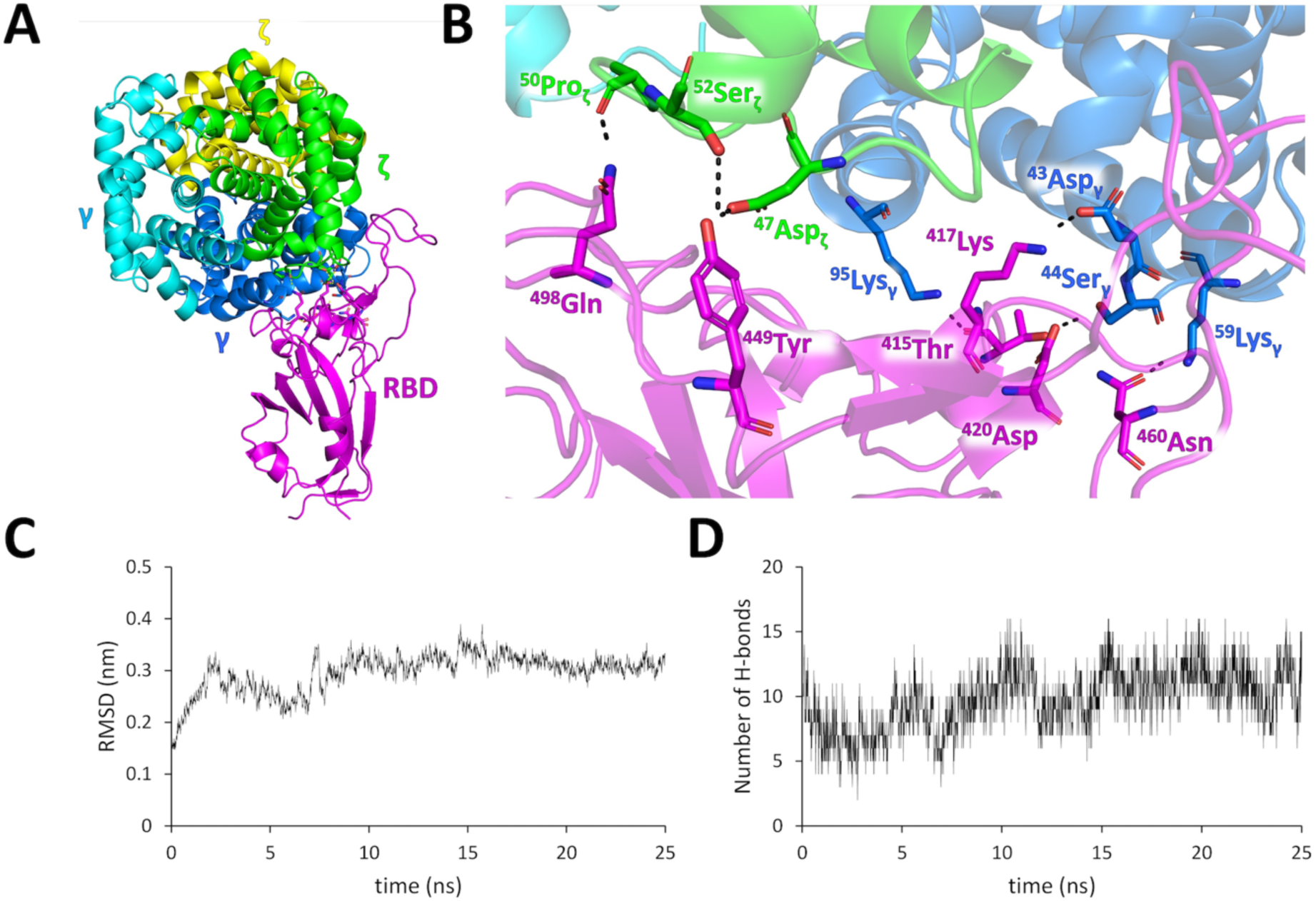
In silico molecular interactions between SARS-CoV-2 Spike and Hb Portland. (A) top scored complex obtained using the HDOCK software. (B) detailed view of hydrogen bonds formed between RBD and Hb Portland. (C) RMSD values obtained during 25 ns of all-atom unbiased molecular dynamics. (D) number of intermolecular hydrogen bonds established during the all-atom molecular dynamics simulation.

### 2.7. The SARS-CoV-2 Spike protein efficiently interacts with Hb Portland: a molecular docking analysis

As reported in other studies, the Spike protein can interact with human hemoglobins [49]. In the case of the K562 system the hemoglobin that is produced at the highest level is Hb Portland (ζ_2_γ_2_) [36]. Therefore, we simulated the interaction between Hb Portland and the S-protein RBD using the well-known protein-protein docking software HDOCK (Figure 7A) [50]. The top scored pose predicted the interaction between the S-protein RBD and both the ζ and γ chains of Portland Hb. Figure 7B shows in detail the H-bonds formed between the proteins. To further strength the reliability of the proposed interaction, the computed model was submitted to 25 ns of all-atom unbiased molecular dynamics simulation. Indeed, the complex remained stable, as it can be seen from the Cα-RMSD values calculated over the simulation time (Figure 7C), with an average number of intermolecular hydrogen bonds equal to 9.6 (Figure 7D). Of note, the hydrogen bonds reported in Figure 7B were retained during the entire molecular dynamics simulation.

## 3. Discussion

The impact of SARS-CoV-2 Spike protein on cellular functions is of key interest, as the two mRNA vaccines BNT162b2 from Pfizer-BioNTech and mRNA-1273 from Moderna, generate high levels of this protein [14, 22–24]. Therefore, searching for circulating Spike in plasma of COVID-19 patient might help in understanding unexpected adverse effects following COVID-19 mRNA vaccination [8, 12]. For instance, Yonker and colleagues were able to identify circulating Spike Protein in patients with Post-COVID-19 mRNA Vaccine myocarditis [18]. Persistent circulating SARS-CoV-2 Spike was recently proposed to be causative of the COVID-19 associated syndrome termed PASC (post-acute sequelae of COVID-19) [25,26]. Considering that the anti-SARS vaccination campaigns are expected to be still ongoing for the next coming years [5], extensive analysis of SARS-CoV-2 Spike in ex vivo cellular systems is required for understanding possible impacts on vaccination [7–9].

The major conclusion of our study is that the BNT162b2 vaccine efficiently transfers the SARS-CoV-2 S-protein mRNA to K562 cells, causing, as expected, production of the S-protein. This was found to be associated with suppression of erythroid differentiation and, more importantly, with sharp inhibition of endogenous and mithramycin induced expression of embryo-fetal globin genes. This was confirmed using different, but convergent, assays (benzidine-staining, RT-qPCR, Western blotting).

In our opinion, the results of this study are of interest when considered together with recently published reports demonstrating that the S-protein has an impact on biological functions of hematopoietic cells [28–30]. In particular, Estep et al. found that SARS-CoV-2 infection and COVID-19 vaccination dramatically impair the functionalities and survivability of hematopoietic stem progenitor cells (HSPCs) in the umbilical cord blood [30]. Collectively these studies suggest that SARS-CoV-2 S-protein, COVID-19 mRNA vaccines and SARS-CoV-2 infection might have dramatic effects of the hematopoietic compartment.

Our data sustain this concept and would stimulate research on the in vivo effects of SARS-CoV-2 infection and/or vaccination not only in healthy subjects, but also in patients affected by hemoglobinopathies. Our study should encourage further investigations on other experimental model systems mimicking erythropoiesis, such as the HUDEP-1 [51] and HUDEP-2 [52] cell lines and, even more importantly, primary erythroid cells isolated from normal subjects and/or patients affected by hemoglobinopathies [53].

In this respect, all the clinical trials on β-thalassemia patients at present ongoing are expected to involve patients vaccinated against SARS-CoV-2. It would be of great interest to compare hematopoietic parameters in these patients with those obtained in similar trials conducted before the COVID-19 pandemic, when the enrolled patients were not vaccinated. Finally, our data should encourage, in our opinion, transcriptomic and proteomic studies to verify the impact of Spike-producing vaccines (such the BNT162b2 from Pfizer-BioNTech [14] and the mRNA-1273 from Moderna [15]) or erythroid pathways.

## 4. Materials and methods

### 4.1. Cell proliferation analysis and erythroid differentiation of K562 cells

Human erythroleukemia K562 cells [31] were seeded at 40,000 cells/mL concentration and the treatments were carried out by adding the appropriate drug concentration as indicated. The proliferation rate (cells number/mL) was analyzed using a model Z2 Coulter counter (Coulter Electronics, Hialeah, FL) after 5 days in order to determine possible effects on cell proliferation. Erythroid differentiation was assessed by benzidine staining and counting blue colored positive cells (percentage of blue cells on 100 cells counted); active benzidine solution was prepared with 0.2% benzidine in 5 M glacial acetic acid adding 10% of the total volume of H_2_O_2_ as described [54].

### 4.2. Treatment with BNT162b2 vaccine and fetal hemoglobin inducer

The BNT162b2 vaccine (COMIRNATY^TM^, Lot. FP8191) was obtained from the Hospital Pharmacy of University of Padova. For treatment with the BNT162b2 vaccine, K562 cells were seeded at 40,000 cells/mL concentration and subsequently treated with increasing concentration of the vaccine (0,5-1-2 μg/mL concentration). After 24 h of treatment, cells were additionally treated with MTH 15 nM in order to induce erythroid differentiation in K562 cells pre-treated with increasing concentration of the vaccine or in K562 cells control cells not treated with vaccine the day before.

### 4.3. RNA extraction from K562 cells

The cells were isolated after 5 days of treatment with MTH to induce erythroid differentiation by centrifugation at 1200 rpm for 8 min at room temperature and lysed in Tri-reagent^TM^ (Sigma-Aldrich, St. Louis, Missouri, USA) following manufacturer’s instruction. The homogenate was incubated for 5 min at room temperature, added with 0.2 mL of chloroform per mL of Tri-reagent^TM^ and vigorously shaken for 15 s, incubated 5 min at room temperature and finally centrifuged at 12,000 rpm for 15 min at 4°C. The aqueous phase was removed and added with 0.5 mL of isopropanol per mL of Tri-reagent^TM^. After 10 min at room temperature, the samples were centrifuged at 12,000 rpm for 15 min at 4 °C. The RNA pellets were washed with 1 mL of 75% ethanol and centrifuged at 12,000 rpm for 10 min at 4 °C. Finally, EtOH was removed and RNA pellets were suspended in Nuclease-free water to proceed with downstream analysis.

### 4.4. RT-qPCR analysis

For the synthesis of cDNA with random hexamers (PrimeScript RT reagent kit from Takara Bio) 300 ng of total RNA were used. Quantitative real-time PCR assay was carried out using gene-specific fluorescently labelled probes and using CFX96 PCR system by Bio-Rad. The nucleotide sequences used for real-time qPCR analysis are showed in Table 1; human glyceraldehyde-3-phosphate dehydrogenase (GAPDH), RPL13A and β-actin were used as reference genes [40].Each reaction mixture contained 1x TaKaRa Ex Taq® DNA Polymerase (Takara Bio Inc., Shiga, Japan), 300 nM forward and reverse primers and the 200 nM probes (Integrated DNA Technologies, Castenaso, Italy). SARS-CoV-2 quantification was performed employing the PowerUp SYBR Green Master Mix (Thermo Fisher Scientific, Inc.) with indicated primers (Table 1). The assays were carried out using CFX96 Touch Real-Time PCR System (Bio-Rad, Hercules, California, USA). After an initial denaturation at 95°C for 1 min, the reactions were performed for 50 cycles (95°C for 15 sec, 60°C for 60 sec). Data were analyzed by employing the CFX manager software (Bio-Rad, Hercules, California, USA). To compare gene expression of each template amplified, the ΔΔCt method was used [55].

**Table 1.**
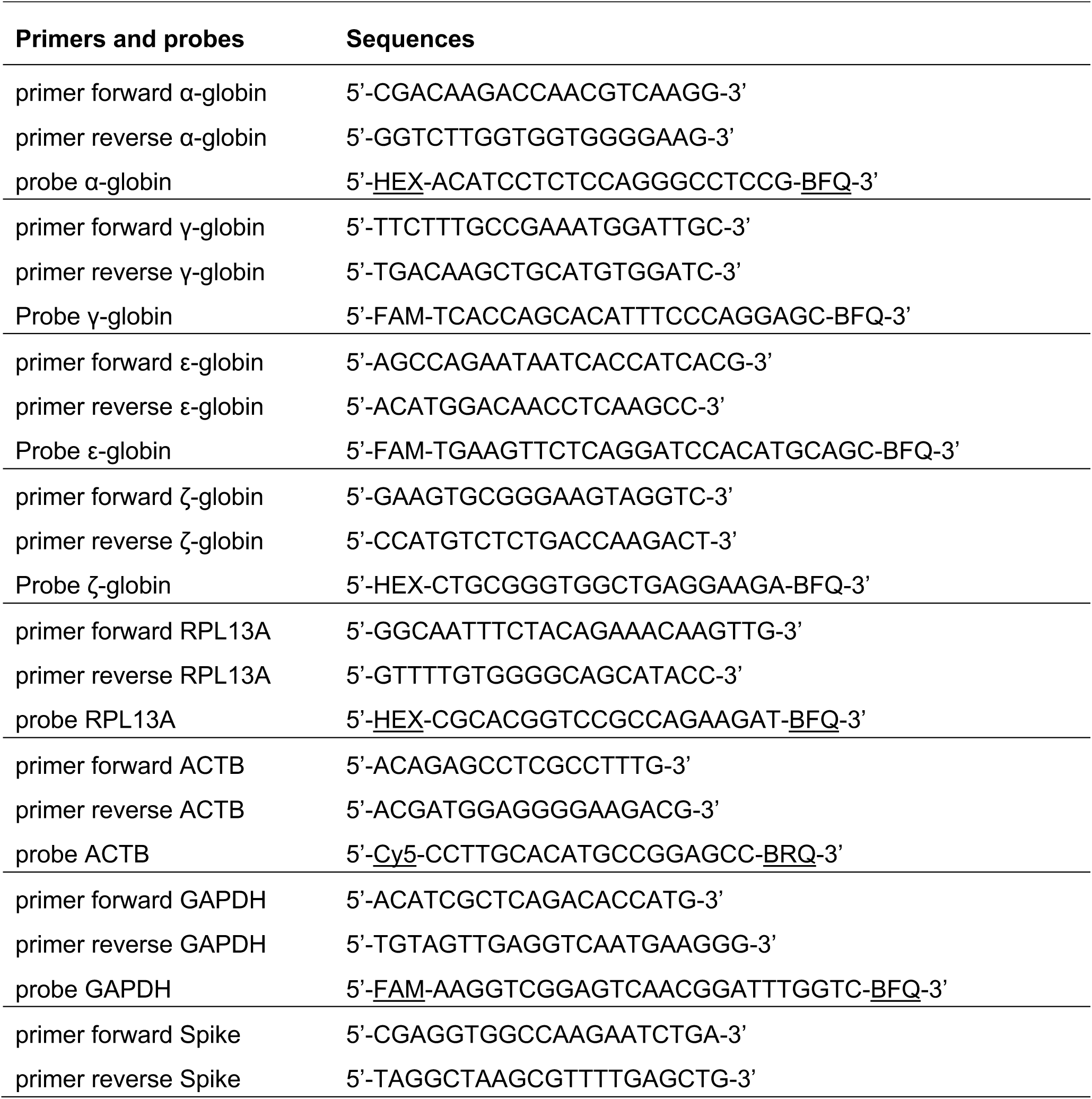
List of primers and probes with related sequences used to perform RT-qPCR analyses on K562 cells.

### 4.5. Western blotting analysis

The accumulation of γ, ζ, and ε-globin proteins (14 kDa) in uninduced or MTH-induced K562 cells cultured in the absence or in the presence of BNT162b2 was assessed by Western blotting. For whole-cell extract preparation, the cells were lysed with RIPA buffer (Thermo Fisher Scientific) following manufacturer’s instruction and quantified by BCA assay (PierceTM BCA Protein Assay kit, Thermo Fisher Scientific). For each sample 20 μg of K562 cell extracts were loaded on 6-18% hand-casted acrylamide SDS-PAGE gradient gel (40% Acrylamide/bis-Acrylamide solution, BioRad). After separation by electrophoretic run, the proteins were transferred onto 0.2 μm nitrocellulose paper (Protran®, Cytiva^TM^), and incubated with the primary antibodies listed in Table 2; the constitutive protein β-Actin was selected as housekeeping to normalize the quantification of the target proteins. Membranes were incubated with an appropriate HRP-conjugated secondary antibody (Cell signalling technologies, cat. n. 7074) and LumiGLO® ECL kit (Cell Signaling Technology) was employed following manufacturer’s instruction before to exposure to X-ray film (Cytiva^TM^). As necessary, after stripping procedure using the Restore™ Western Blot Stripping Buffer (Thermo Fisher Scientific) membranes were re-probed with primary and secondary antibodies as previously described [56]. The quantification of obtained bands was carried out by ChemiDoc (Bio-Rad) and densitometric analysis was performed with Image Lab Software (Bio-Rad).

**Table 2.** List of primary antibodies used to perform Western Blot analysis with their manufacturers.

| **Target** | **Primary antibody** | **Cat.n.** |
| --- | --- | --- |
| γ-globin | Rabbit anti-γ-globin<br>(Thermo Fisher Scientific Inc., Waltham, MA, USA) | PA5-29006 |
| ζ-globin | Rabbit anti-ζ-globin<br>(ABclonal, Woburn, MA, USA) | A6920 |
| ε-globin | Rabbit anti-ε-globin<br>(ABclonal, Woburn, MA, USA) | A3909 |
| Transferrin<br>receptor (CD71) | Rabbit anti-TfR<br>(ABclonal, Woburn, MA, USA) | A22161 |
| β-actin | Rabbit anti-β-actin<br>(Cell Signalling Technology, Danvers, MA, USA) | 4967 |

### 4.6. Computational studies

All the computational methodologies were carried out on a 32 Core AMD Ryzen 93,905×, 3.5 GHz Linux Workstation (O.S. Ubuntu 20.04) equipped with GPU (Nvidia Quadro RTX 4000, 8 GB). The SARS-CoV-2 Spike receptor binding domain (RBD) was retrieved from the Protein Data Bank (PDB-ID: 7kn5). The Hb Portland structure was obtained by replacing the α chains of a fetal hemoglobin structure (α_2_γ_2_; PDB-ID: 4mqj) with the ζ chains of the available ζ_2_β_2_ hemoglobin crystallographic structure (PDB-ID: 3w4u). HDOCKlite v1.0 software [50] was then used to predict the interaction geometry between the proteins and the top scored complex was submitted to all-atom unbiased molecular dynamics (MDs) simulation using the GROMACS software [57] patched with the open-source, community developed Plumed ver 2.6.5 [58] under the Charmm36 force field [59]. The complex was included in a rectangular box of 10 x 10 x 15 nanometers length, solvated and neutralized using 0.15M potassium chloride. The full system was submitted to energy minimization and equilibrated under NVT and NPT conditions. Long range electrostatic interactions were modelled using the Particle Mesh Ewald algorithm. LINCS, Nosé-Hoover and Parrinello-Rahman algorithms were used in the simulations for restraints, and as thermostat and barostat respectively. MDs were conducted under the NPT conditions for 25 ns with 2 fs time steps. Root-mean-squared deviation (RMSD) and number of hydrogen bonds were obtained through the “rms” and “hbond” tools implemented in Gromacs.

### 4.7 Statistics

All the data were normally distributed and presented, unless otherwise stated, as mean ± S.D. Statistical differences between groups were compared using one-way ANOVA (analyses of variance between groups) followed by Dunnett’s multiple comparison or paired t-test employing Prism (v. 9.02) by GraphPad software. Statistical differences were considered significant when *p* < 0.05 (*), and highly significant when *p* < 0.01 (**) and p < 0.001 (***).

## Credit author statement

Matteo Zurlo: Investigation, Formal analysis, Writing – original draft. Jessica Gasparello: Investigation. Marco Verona: Investigation. Chiara Papi: Investigation. Lucia Carmela Cosenza: Investigation. Alessia Finotti: Supervision, Funding acquisition, Writing-Reviewing and Editing. Giovanni Marzaro: Supervision, Funding acquisition, Writing-Reviewing and Editing. Roberto Gambari: Conceptualization, Supervision, Funding acquisition, Writing-Reviewing and Editing.

## Data availability

The datasets used and/or analyzed during the current study are available from the corresponding author on reasonable request.

## Supporting information

Surlo-SUPPLEMENTARY

## Acknowledgments

We thank “Associazione tutti per Chiara” (Montagnana, Italy) for supporting M.Z., C.P. and L.C.C. with fellowships.

## Appendix A

Supplementary data

