## Supplementary material for "The anti-SARS-CoV-2 BNT162b2 vaccine suppresses mithramycin-induced erythroid differentiation and expression of embryo-fetal globin genes in human erythroleukemia K562 cells": Surlo-SUPPLEMENTARY: Supplementary-Figure-S1.docx

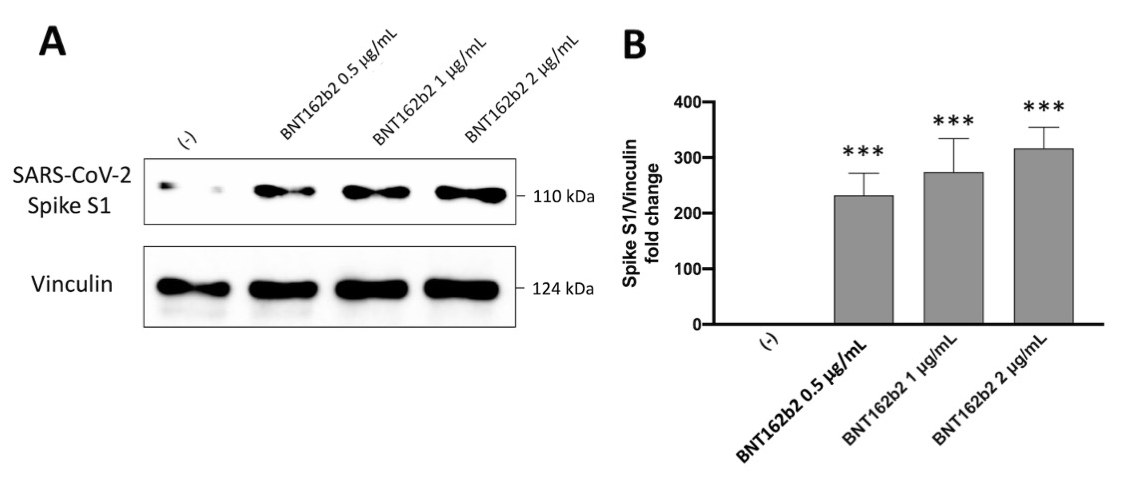

**Figure S1. SARS-CoV-2 Spike protein is produced by K562 cells treated with the BNT162b2 vaccine.** K562 cells were treated for 6 days with 0.5, 1 and 2 μg/mL of the BNT162b2 vaccine. The accumulation of S-protein (approximately 110 kDa) in K562 cells cultured in the absence or in the presence of BNT162b2 was assessed by Western blotting. For whole-cell extract preparation, the cells were lysed with RIPA buffer (Thermo Fisher Scientific) following manufacturer’s instruction and quantified by BCA assay (PierceTM BCA Protein Assay kit, Thermo Fisher Scientific). For each sample 20 μg of K562 cell extracts were loaded on 7% hand-casted acrylamide SDS-PAGE gel (40% Acrylamide/bis-Acrylamide solution, BioRad). After separation by electrophoretic run, the proteins were transferred onto 0.2 μm nitrocellulose paper (Protran®, Cytiva^TM^), and incubated with the primary antibody against SARS-CoV-2 S1 subunit (A20834, ABclonal, Woburn, MA, USA); the constitutive protein Vinculin (CSB-PA13779A0Rb, Cusabio, Houston, TX, USA) was selected as housekeeping to normalize the quantification of the target protein. Membranes were incubated with an appropriate HRP-conjugated secondary antibody (Cell signalling technologies, cat. n. 7074) and LumiGLO® ECL kit (Cell Signaling Technology) was employed following manufacturer’s instruction before to exposure to X-ray film (Cytiva^TM^). As necessary, after stripping procedure using the Restore™ Western Blot Stripping Buffer (Thermo Fisher Scientific) membranes were re-probed with primary and secondary antibodies. The quantification of obtained bands was carried out by ChemiDoc (Bio-Rad) and densitometric analysis was performed with Image Lab Software (Bio-Rad). Results are presented as mean ± S.E.M; statistical differences between groups were compared using ANOVA. (***): p < 0.001 (highly significant). The S-protein mRNA content following treatment of K562 cells with BNT162b2 is shown in Figure 1B of the main text (RT-qPCR analysis).
