## Supplementary material for "The anti-SARS-CoV-2 BNT162b2 vaccine suppresses mithramycin-induced erythroid differentiation and expression of embryo-fetal globin genes in human erythroleukemia K562 cells": Surlo-SUPPLEMENTARY: Supplementary-Figure-S2.docx

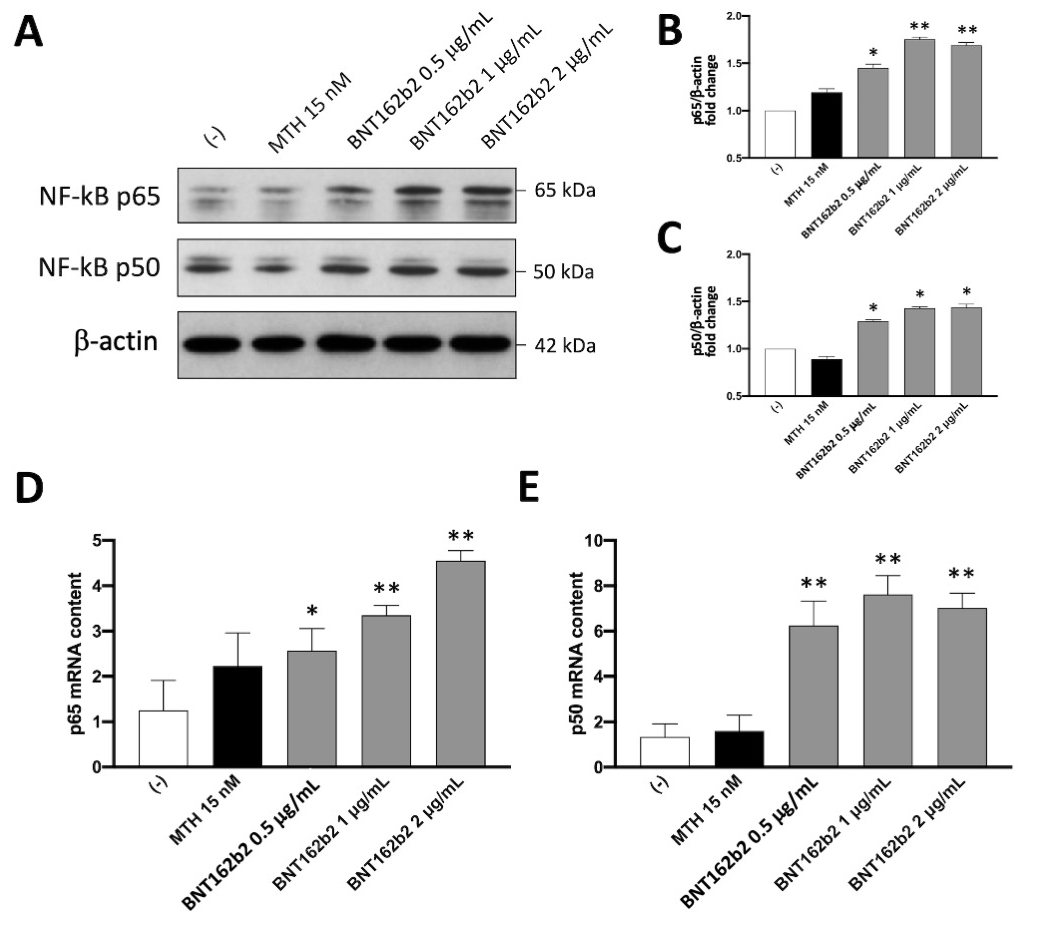

**Figure S2. Expression of NF-kB protein in K562 cells treated with the BNT162b2 vaccine.** K562 cells were treated for 6 days with 0.5, 1 and 2 μg/mL of the BNT162b2 vaccine. The accumulation of NF-kB p50 and p65 (approximately 50 and 65 kDa, respectively) in K562 cells cultured in the absence or in the presence of BNT162b2 was assessed by Western blotting. Technical details can be found in the legend to Figure S1. After separation by electrophoretic run, the proteins were transferred onto 0.2 μm nitrocellulose paper (Protran®, Cytiva^TM^), and incubated with the primary antibodies against NF-kB p50 (GTX133711, GeneTex, Irvine, CA, USA) or NF-kB p65 (GTX102090, GeneTex, Irvine, CA, USA) (A); the constitutive protein β-actin (4967, Cell Signalling Technology, Danvers, MA, USA) was selected as housekeeping to normalize the quantification of the target proteins (B and C). Obtained data shows a good correlation also analyzing the mRNA content of p65 and p50 by RT-qPCR (D and E respectively); employed primers and probes are listed in Supplementary Table S1. Results are presented as mean ± S.E.M; statistical differences between groups were compared using ANOVA. (*): p < 0.05 (significant); (**): p < 0.01 (highly significant).
