## Supplementary material for "The anti-SARS-CoV-2 BNT162b2 vaccine suppresses mithramycin-induced erythroid differentiation and expression of embryo-fetal globin genes in human erythroleukemia K562 cells": Surlo-SUPPLEMENTARY: Supplementary-Figure-S3.docx

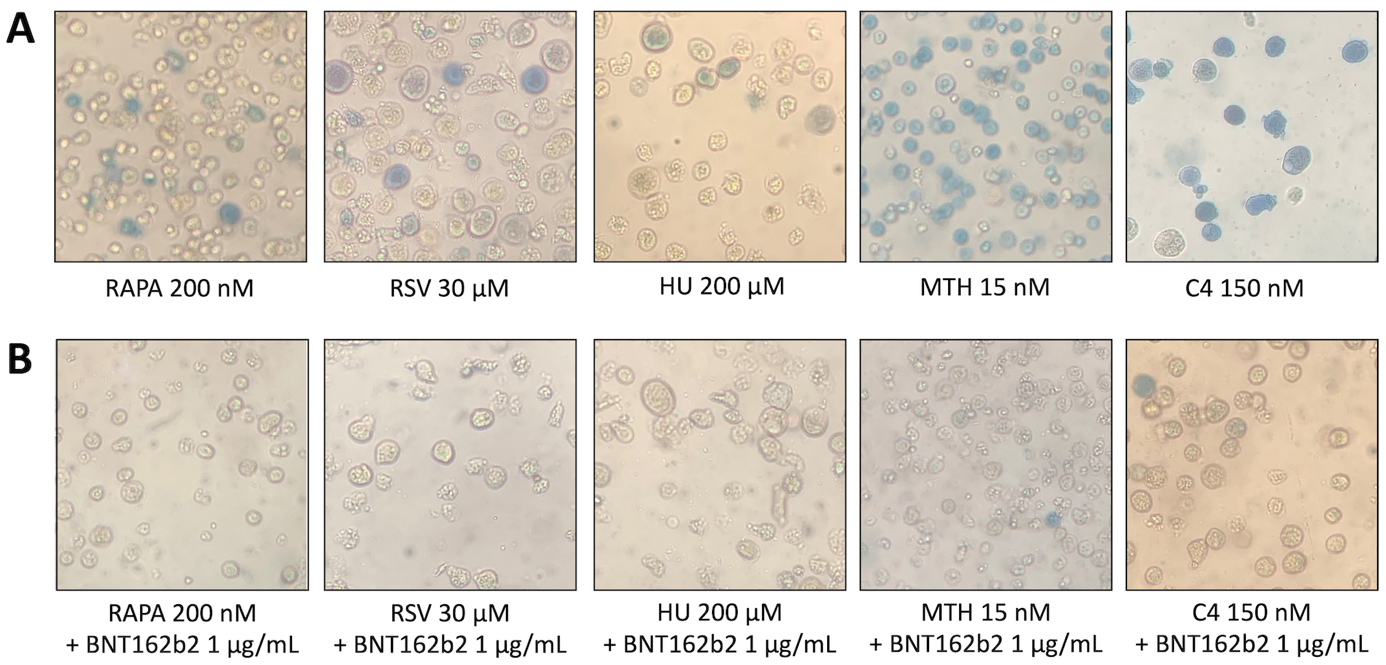

**Figure S3.** Effects of BNT162b2 vaccine on erythroid differentiation induced in K562 cells by different inducers: benzidine-staining. K562 cells were induced with 200 nM rapamycin (RAPA), 30 μM resveratrol (RSV), 200 μM hydroxyurea (HU), 150 nM isoxazole C4 in the absence (A) or in the presence (B) of 1 μg/mL BNR162b2, as indicated. Benzidine assay was performed after 5 days of treatment with hemoglobin inducers; 15 nM Mithramycin (MTH) was used as a positive control.
