## Supplementary material for "The anti-SARS-CoV-2 BNT162b2 vaccine suppresses mithramycin-induced erythroid differentiation and expression of embryo-fetal globin genes in human erythroleukemia K562 cells": Surlo-SUPPLEMENTARY: Supplementary-Table-S1.docx

**Table S1.** List of primers and probes used for NF-kB p50 and p65 detection by RT-qPCR.

| **Primers and probes** | **Sequences** |
| --- | --- |
| primer forward p50 | 5’-GGATCTGCACTGTAACTGCT-3’ |
| primer reverse p50 | 5’-CTCTGTCATTCGTGCTTCCA-3’ |
| probe p50 | 5’-FAM-TGTCACATGAAGTATACCCAGGTTTGCG-BFQ-3’ |
| primer forward p65 | 5’-CGAGCTTGTAGGAAAGGACTG-3’ |
| primer reverse p65 | 5’-TGACTGATAGCCTGCTCCAG-3’ |
| Probe p65 | 5’-FAM-CGCTGCATCCACAGTTTCCAGAAC-BFQ-3’ |
